# Establishing gene regulatory networks from Parkinson’s disease risk loci

**DOI:** 10.1101/2021.04.08.439080

**Authors:** Sophie L. Farrow, William Schierding, Sreemol Gokuladhas, Evgeniia Golovina, Tayaza Fadason, Antony A. Cooper, Justin M. O’Sullivan

## Abstract

The latest meta-analysis of genome wide association studies (GWAS) identified 90 independent single nucleotide polymorphisms (SNPs) across 78 genomic regions associated with Parkinson’s disease (PD), yet the mechanisms by which these variants influence the development of the disease remains largely elusive. To establish the functional gene regulatory networks associated with PD-SNPs, we utilised an approach combining spatial (chromosomal conformation capture) and functional (expression quantitative trait loci; eQTL) data. We identified 518 genes subject to regulation by 76 PD-SNPs across 49 tissues, that encompass 36 peripheral and 13 CNS tissues. Notably, one third of these genes were regulated via *trans*-acting mechanisms (distal; risk locus-gene separated by > 1Mb, or on different chromosomes). Of particular interest is the identification of a novel *trans*-eQTL-gene connection between rs10847864 and *SYNJ1* in the adult brain cortex, highlighting a convergence between familial studies and PD GWAS loci for *SYNJ1 (PARK20)* for the first time. Furthermore, we identified 16 neuro-development specific eQTL-gene regulatory connections within the foetal cortex, consistent with hypotheses suggesting a neurodevelopmental involvement in the pathogenesis of PD. Through utilising Louvain clustering we extracted nine significant and highly intra-connected clusters within the entire gene regulatory network. The nine clusters are enriched for specific biological processes and pathways, some of which have not previously been associated with PD. Together, our results not only contribute to an overall understanding of the mechanisms and impact of specific combinations of PD-SNPs, but also highlight the potential impact gene regulatory networks may have when elucidating aetiological subtypes of PD.

## Introduction

Parkinson’s disease (PD) is considered to be primarily an idiopathic neurodegenerative disorder, with monogenic forms contributing to just 5-10% of all cases^1^. However, the idiopathic nature of PD is being questioned, as evidence increasingly supports a complex involvement of genetics in the development of the majority of cases^2,3^. Genome-wide association studies (GWAS) have identified >200 PD risk loci^4^, with 90 PD-associated Single Nucleotide Polymorphisms (PD-SNPs) across 78 risk loci reported in the largest meta-analysis to date^5^. As is typically observed with GWAS variants, the majority of the PD-SNPs are located within non-coding regions of the genome, with no direct or obvious influence on protein structure or function^6,7^. Studies have shown that such non-coding disease-associated variants are more likely to be located within regulatory regions^8^, and thus contribute to risk through influencing gene regulation and expression, either locally or distally. These regulatory interactions are likely to be tissue specific, adding a further layer of complexity. Consequently, determining how these variants contribute to PD risk, both individually and in combination, poses a major scientific challenge^9,10^.

Although PD is defined as a neurodegenerative disease, mounting evidence demonstrates the role of non-central nervous system (CNS) tissues in the development and presentation of such disorders (*i.e*. Huntington’s^11^ and PD^12,13^). Both alpha-synuclein (αSyn) protein pathology and modulation of PD-related genes have been identified in peripheral tissues (*e.g*. the gastrointestinal tract and heart) of PD patients^12,14–19^. The contribution of peripheral tissue in the origins of PD warrants further research, and thus the consideration of how PD-SNPs mediate risk should not be confined to tissues of the CNS.

Spatial gene regulatory interactions are hypothesized to be drivers of complex trait heritability^20^, acting through both *cis*-(nearby) and *trans*-(distal; locus-gene separated by > 1Mb, or on different chromosomes) mechanisms (Figure 1)^19,21,22^. These *cis*- and *trans*-acting elements can regulate the transcription of one or more genes, in a tissue specific manner, and are commonly detected in the form of expression quantitative trait loci (eQTL)^23^. Genetic variation within elements of gene regulatory networks likely confer risk at different developmental stages, including during foetal neurodevelopment – a critical stage that has a growing body of support in neurodegenerative diseases^24^.

**Figure 1:**
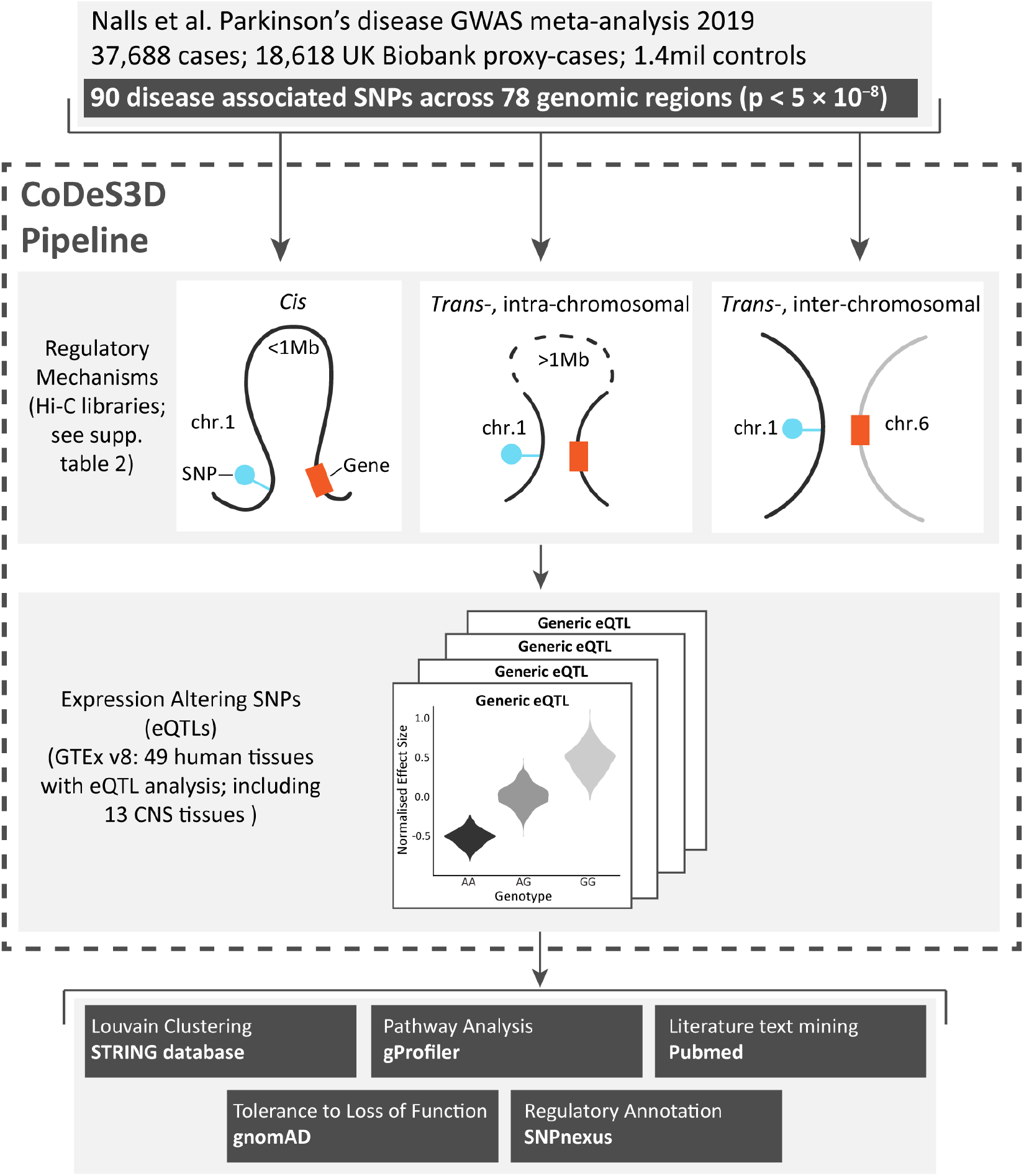
Methods workflow. 90 PD-SNPs were obtained from Nalls et al^5^. Spatial interactions between the 90 PD-SNPs and genes were identified from Hi-C libraries (Supplementary table 2). The resulting spatial SNP-gene pairs were then used to query GTEx v8 to identify significant eQTLs. The resulting SNP-gene pairs were then analysed for functional relevance using multiple tools and databases (methods). Figure adapted from Schierding et al.^19^

Here we performed correlational analyses of experimentally derived data to identify eQTLs that physically connect PD-SNPs to the genes that they control, in three-dimensions, with the goal of understanding the putative functional impacts of known PD-SNPs^21^. The integration of spatial and eQTL data allows for the identification of *trans*-eQTL-gene associations^25^, thereby nominating genes which have not previously been implicated in PD. Our analysis identified 518 genes subject to regulation by 76 PD-SNPs across 49 tissues. Further, clustering analysis of the entire gene network revealed nine significant, intra-connected clusters, enriched for both novel and known PD biological pathways, highlighting putative disease-causative molecular mechanisms and areas for future research.

## Methods

### Data and reference files

The 90 PD-SNPs (across 78 genomic regions; Supplementary table 1) investigated in this study were previously identified by a GWAS meta-analysis as being of genome wide significance (p<5×10^−8^)^5^. The human genome reference sequence used in this study was GRCh38 (hg38).

### Identification of eQTL-gene pairs

The contextualise developmental SNPs using 3D information (CoDeS3D)^21^ algorithm was used to identify genes whose transcript levels are putatively regulated by the 90 PD-SNPs. CoDeS3D integrates data on spatial interactions between genomic loci (Hi-C data; Supplementary table 2) with expression data (genotype-tissue expression database version 8; GTEx v8^26^) to identify genes whose transcript levels are associated with a physical connection to the SNP (*i.e*. spatial eQTL).

Hi-C captures regions of the genome that are physically interacting, and can be covalently connected by a cross-linking agent^27^. The hg38 reference genome was digitally digested with MboI, DpnII and HindIII to obtain all possible Hi-C fragment locations for the 90 PD-SNP loci. All identified SNP fragments (tagged by the PD-SNPs) were then queried against the Hi-C databases (70 different cell lines from 12 studies; Supplementary table 2) to identify distal fragments of DNA that spatially connect to the SNP loci. Spatial SNP-gene connections are established when the SNP-containing fragment spatially connects to a fragment that overlaps any region between the start and end of a gene as defined by GENCODE. There was no binning or padding around restriction fragments to obtain gene overlap. The resulting spatial SNP-gene pairs were subsequently used to query the GTEx v8 eQTL database^26^ to identify spatial SNP-gene pairs with significant eQTLs (both *cis*- and *trans*-acting eQTL; false discovery rate (FDR) adjusted *p* < 0.05).

We performed a ‘brain specific’ analysis by interrogating only the subset of Hi-C libraries derived from brain-specific cell lines (11 cell lines from 4 studies, highlighted in red in Supplementary table 2)^28–31^ and only expression data from the 13 brain-specific tissues in GTEx v8.

### Identification of neurodevelopmental-specific eQTL-gene pairs

We performed a neurodevelopmental stage specific analysis by interrogating Hi-C libraries from foetal-specific brain cell lines (cortical plate neurons; germinal zone neurons; Supplementary table 2; datasets no. 1 and 2) with expression data from a foetal cortex eQTL dataset^32^.

### Functional analysis of eQTL SNPs

SNPnexus v4^33^ (https://www.snp-nexus.org/v4/; accessed 22/06/2020) was used to obtain known epigenomic annotations for the eQTL SNPs.

### Probability of gene loss of function intolerance

The LOEUF (loss-of-function observed/expected upper bound fraction) ^34^ score for the genes, within the significant SNP-gene pairs, was obtained from gnomAD v2.1.1^35^ (https://gnomad.broadinstitute.org/; accessed 21/07/2020) to determine the level of constraint on the identified genes.

### Protein-protein interaction (PPI) and Modularity clustering

STRING^36^ (Search Tool for Retrieval of Interaction Genes/proteins; https://string-db.org, accessed 22/07/2020) was queried to identify published information on interactions between genes, or their respective proteins. Only PPIs with a high confidence level (>0.700, as defined in STRING) were used for this analysis. A Louvain method was used to determine the syntality of each node, following four different criteria: 1) immediate connection; 2) shortest path (*i.e*. the minimum number of edges connecting any two nodes); 3) node acting as a bridge; 4) connections that nodes have in common. The proteins were then hierarchically clustered using the Louvain algorithm^37^, clusters were defined as significant if p < 0.05.

### Pathway analyses & literature search

The g:Profiler^38^ database was used to identify enriched pathways. Queries were run on: 1) all genes’ 2) the ‘*cis*’; and 3) the ‘*trans*’ subsets of genes (*i.e*. genes regulated only in *cis*, or only in *trans*). PubMed was used to search (using “[gene name]” and Parkinson’s”) the published literature for all identified genes and known PD associations.

### Additional analyses of genes and variants identified by *Makarious* et al

*Makarious* et al recently utilised a multi-modality approach to identify genetic and transcriptomic features that contribute to risk predictions of PD^39^. They highlighted two SNPs (rs10835060 and rs4238361), and 29 genes. We performed CoDeS3D analysis on the two SNPs, across all Hi-C cell lines and GTEx tissues (as previously described). The resulting eQTL gene pairs, along with the 29 genes highlighted through transcriptomic analysis, were combined with our set of 523 genes. Louvain clustering and PPI analysis were re-run on this combined list of genes to see how or if the subset of genes co-locate within the networks.

### URLs

CoDeS3D pipeline: https://github.com/Genome3d/codes3d-v2

gnomAD: https://gnomad.broadinstitute.org/

gProfiler: https://biit.cs.ut.ee/gprofiler/

UCSC: https://genome.ucsc.edu/index.html

STRING: https://string-db.org/

SNPNexus: https://www.snp-nexus.org/

Louvain clustering analysis: https://github.com/Genome3d/PPI-network-analysis

### Data Availability

All data generated during this study are included in the supplementary information. Datasets analysed and tools used in this study were all derived from publicly available resources (See URLs).

## Results

### PD GWAS SNPs regulate the expression of > 500 genes

*Nalls* et al.^5^ identified 90 SNPs that were associated with PD at the level of genome-wide significance (Supplementary table 1), yet the mechanisms by which these variants influence the development of the disease remains largely elusive. We used the CoDeS3D algorithm^21^ to identify SNPs that have evidence of physical interaction with the gene as captured by Hi-C (Supplementary table 2) and also associate with changes in gene expression (hereafter eQTLs) and the genes whose transcript levels were affected (Figure 1).

76 (84%) of the 90 PD risk SNPs were identified as eQTLs associated with the regulation of 518 genes through 542 unique eQTL-gene pairs, across the 49 tissues (Table 1, Supplementary table 3). The 76 eQTLs were individually associated with the regulatory impacts of as few as one, or as many as 39 genes in *cis* and *trans* (Supplementary figure 1). We identified 178 of the 542 genes as being associated with PD through *trans*-eQTL-gene connections. We did not identify eQTL interactions for 14 of the 90 SNPs. Conversely, 8 of these 14 SNPs are annotated as being eQTLs in the IPDGC GWAS Locus Browser^40^, and a further four are eQTLs in GTEx (Supplementary table 1). However, in these 12 instances, the eQTLs occur in *cis*-(*i.e*. within 1Mb) and are not supported by Hi-C data.

**Table 1:** Summary statistics for the spatial eQTL-gene regulatory network for the 90 PD-SNPs. SNPs were downloaded from Nalls et al 2019 GWAS paper (download date: 18.06.2020).

<sup>#</sup>eQTL SNPs were defined as having significant spatial interactions (FDR
|  | PD SNPs |  |
| --- | --- | --- |
|  | Brain specific* | All tissues* |
| No. SNPs | 90 | 90 |
| No. eQTL SNPs <sup>#</sup> | 55 | 76 |
| No. Genes <sup>†</sup> | 165 | 518 |
| No. eQTL-Gene pairs <sup>‡</sup> | 167 | 542 |
| No. *trans* eQTL-Gene pairs | 30 | 178 |

$\leq 0.05$ ) with at least one gene. <sup>¶</sup>genes were those whose expression was shown to be affected by an eQTL SNP. <sup>‡</sup> The total number of SNP-gene pairs reflects interactions with $FDR \leq 0.05$ in at least one GTEx tissue. \* For full list of tissues see Supplementary tables 3 and 4.

Consistent with observations for SNPs associated with other traits^41^, at least one *trans*-regulatory interaction was identified for 81.6% (62 of 76) of the eQTLs. Moreover, 92.7% (165 of 178) of these *trans*-eQTL-gene interactions were identified in only one tissue. By contrast, the *cis*-interactions were identified in eight tissues on average (range of 1 to 49 tissues). 11.8% of the eQTLs (9 of 76; Supplementary table 1) were exclusively involved in *trans*-regulatory interactions. *Trans*-eQTL interactions regulated 18.1% of the genes identified in the brain (30 out of 166; Supplementary table 4), and 32.8% of the genes amongst all 49 tissues (178 out of 518; Table 2; Supplementary table 3). Collectively, these results highlight the importance of looking beyond the nearest gene to identify the regulatory effects of disease-associated variants.

**Table 2:** Proportion of genes subject to cis- and trans-regulation. The proportion of eQTL-gene pairs that are either cis-, trans-intrachromosomal or trans-interchromosomal in (a) 13 GTEx brain-specific tissues and (b) all 49 GTEx tissues. Brain-specific indicates the eQTL dataset obtained through analysing Hi-C cell lines only from the brain, and eQTLs only from the brain tissues in GTEx. All-tissues indicates the eQTL dataset obtained through analysing all Hi-C cell lines, and eQTLs from all tissues in GTEx. There is a significant difference (Chi square test p.value < 0.01) between brain tissues and all tissues for the proportions of the cis vs. trans eQTLs.

|  | Genes subject to *cis*- or *trans*- regulation |  |
| --- | --- | --- |
|  | Brain specific* | All tissues* |
| *Cis* eQTL-gene pairs | 136 (82.0%) | 364 (67.2%) |
| *Trans*-intrachromosomal eQTL-gene pairs | 10 (6.0%) | 56 (10.3%) |
| *Trans*-interchromosomal eQTL-gene pairs | 20 (12.0%) | 122 (22.5%) |

We reasoned that SNPs that are involved in eQTLs likely mark enhancer or promoter sites^42^. We queried SNPnexus^33^ to identify those eQTLs that were marked by histone modifications or fell within open chromatin regions, as indicated by DNAse accessibility. Consistent with our hypothesis, 91% (69 of 76) of the SNPs were marked by histone modifications associated with either enhancers (58) and/or promoters (27). 27.6% of the SNPs were within accessible chromatin(Supplementary table 5). Collectively, these results are consistent with the hypothesis that the loci marked by these eQTLs may be involved in the regulation of gene expression.

Pathway analysis was conducted on the complete set of 518 genes that were impacted by the eQTLs (Supplementary table 6). g:Profiler^38^ identified significant (adj.p < 0.05) enrichment within 10 known biological pathways (g:GOSt), including response to interferon-gamma, synaptic vesicle recycling and endocytosis.

### The regulatory impact of PD-SNPs extends beyond the CNS

Although PD is considered a degenerative disease of the brain, it has become apparent that dysfunction and or alpha-synuclein pathology is observed in non-CNS tissues of PD patients^13–15^. Our spatial eQTL analysis included an assessment of the tissue distribution of the effects of the identified eQTLs within 13 CNS and 36 peripheral tissues. We identified peripheral tissue-specific eQTLs for 28% of the PD-SNPs (21 of 76). Only 2 of the 76 PD-SNPs (i.e. rs10756907 – *SH3GL2*, brain cortex; rs873786 – *SLC26A1*, brain cerebellum) had eQTLs that impacted gene expression levels exclusively in the brain. This supports a possible role for peripheral tissues in PD risk (Supplementary figure 3).

The ability to detect eQTLs in specific tissues is known to correlate with tissue sample size within GTEx^43^. Consistent with this, we identified highly significant correlations between tissue sample numbers and a) all-eQTLs in the brain tissues (Figure 2a; identified using brain specific Hi-C and eQTL data; Supplementary Table 4); and b) all tissues (i.e. the 49 tissues included within GTEx; Figure 2e). These highly significant correlations remained when analysing the *cis*-eQTL subsets in the brain (R = 0.93, p = 3.6e^-06^; Figure 2b), and all tissues (R = 0.9, p < 2.2e^-16^; Figure 2f). Similarly, the correlation was evident for *trans*-intrachromosomal eQTLs detected in all tissues (R = 0.67, p < 1.8e^-06^; Figure 2g). By contrast, there was no observable correlation between the number of *trans*-interchromosomal interactions and tissue sample number (Figure 2d,h). The substantia nigra and brain cerebellar hemisphere exhibited more *trans*-interchromosomal-eQTLs (Figure 2d), while the thyroid exhibited more eQTLs than expected across all three categories (Figure 2e-h).

**Figure 2:**
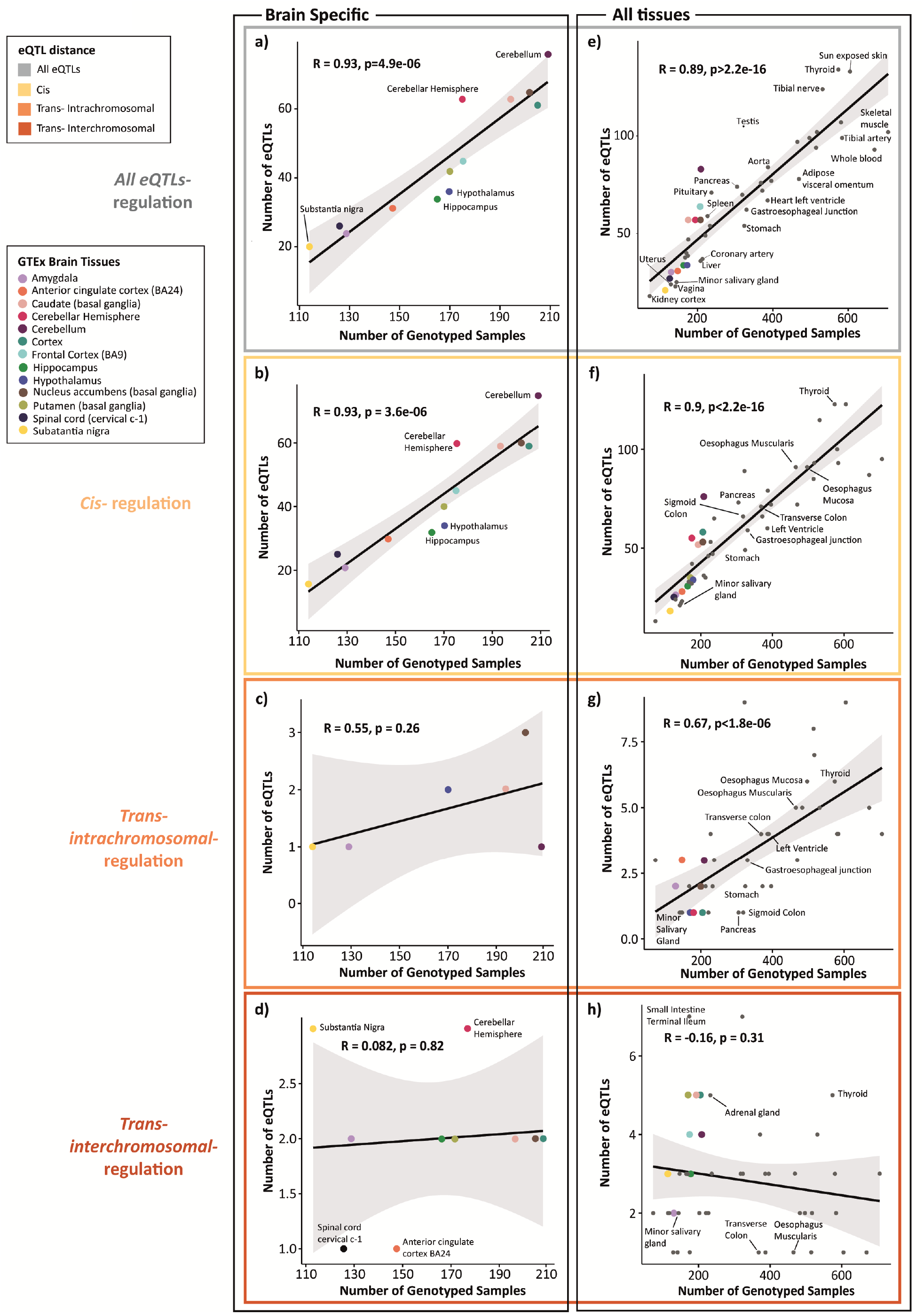
Correlation between genotype samples per tissue and number of eQTLs present in the tissue. (a) Correlation between the number of genotyped samples per tissue (in GTEx) and the number of eQTLs (including cis, trans-intrachromosomal and trans-interchromosomal) per tissue, in 13 brain-specific tissues (b) Correlation between the number of genotyped samples per tissue (in GTEx) and the number of cis-eQTLs per tissue, in 13 brain-specific tissues (c) Correlation between the number of genotyped samples per tissue (in GTEx) and the number of trans-intrachromosomal-eQTLs per tissue, in 13 brain-specific tissues (d) Correlation between the number of genotyped samples per tissue (in GTEx) and the number of trans-interchromosomal-eQTLs per tissue, in 13 brain-specific tissues (e) Correlation between the number of genotyped samples per tissue (in GTEx) and the number of eQTLs (including cis, trans-intrachromosomal and trans-interchromosomal) per tissue, in all 49 tissues (f) Correlation between the number of genotyped samples per tissue (in GTEx) and the number of cis-eQTLs per tissue, in all 49 tissues (g) Correlation between the number of genotyped samples per tissue (in GTEx) and the number of trans-intrachromosomal-eQTLs per tissue, in all 49 tissues (h) Correlation between the number of genotyped samples per tissue (in GTEx) and the number of trans-interchromosomal-eQTLs per tissue, in all 49 tissues. The tissues that fall furthest from the confidence interval are annotated. The grey dots show the correlation for all GTEx tissues. The 13 brain tissues (from GTEx) are indicated by the coloured dots, as shown in the legend. For information on all tissues outside of the 95% CI see Supplementary table 12.

### Genes subject to *trans*-regulation by PD-SNPs are more likely to be intolerant to loss-of-function mutations

Genes that are intolerant to inactivation by loss-of-function variants are deemed essential for healthy development^44^. Intolerance to loss of function variants leaves changes to regulation as one of the few mechanisms that can be modified to introduce variation at a population level. The 117 *trans*-interchromosomal-eQTL regulated genes were significantly (p < 0.01, Kruskal-Wallis test) more intolerant to loss-of-function mutations (LOEUF 0.42 [median]; a low LOEUF score is indicative of evolutionary constraint) than those regulated by *cis*- or *trans*-intrachromosomal acting eQTLs (LOEUF 0.83 and 0.85 respectively [median]; Figure 3; Supplementary table 7). This result is consistent with earlier observations that *trans*-eQTLs are enriched in regulating constrained genes with low LOEUF scores^19^.

**Figure 3:**
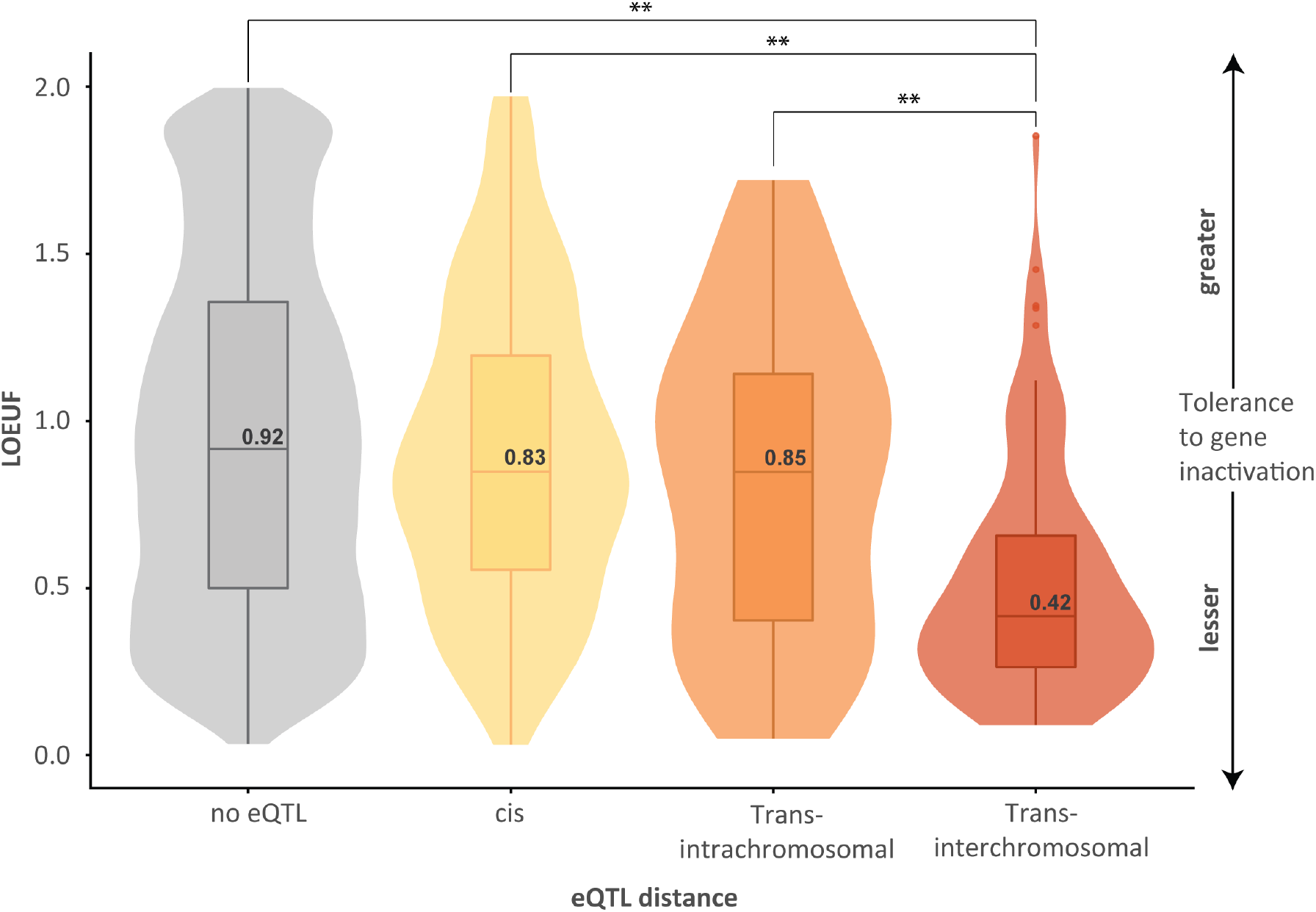
Genes subjected to trans-regulation by PD-SNPs are enriched for Loss-of-Function Intolerance. Genes that are loss of function intolerant, as measured by a continuous LOEUF score, are enriched in *trans*-regulatory interactions involving PD-SNPs. The LOEUF score is a continuous value that indicates the tolerance of a given gene to inactivation. Low LOEUF scores indicate stronger selection against loss-of-function variation. The distribution is shown as a violin plot with the median (LOEUF) values for each eQTL group (black text). The groups were compared using a Kruskal-Wallis test (** = *P* value < 0.01); the absence of a significance value indicates the LOEUF values of the two groups were not significantly different. No eQTL = all genes in gnomAD with an assigned pLI or LOEUF for which an eQTL was not identified in this study (~18,500 genes). Not all genes had LOEUF scores (Supplementary figure 2; Supplementary table 7)

### PD GWAS SNPs regulate expression of a subset of genes within the foetal cortex

Emerging evidence suggests PD has a neuro-developmental aspect^48^, similar to recent observations in Huntington’s disease^24^. Therefore, we analysed the regulatory impacts of the PD-SNPs using foetal cell line Hi-C (i.e. cortical plate neurons and germinal zone neurons^29^) and foetal cortex eQTL datasets^32^ (Supplementary table 8). 33 genes were found to be regulated by 22 PD-SNPs in the foetal cortex. Of these, sixteen genes were regulated by eQTLs involving PD-SNPs in the foetal cortex, without evidence of any eQTLs in adult brain tissues (Figure 4; Supplementary table 4). Ten genes were affected by eQTLs involving PD-SNPs in both the foetal and adult cortex, with effect sizes that were similar in both (Figure 4). Finally, seven genes were regulated by *cis*-eQTLs in the foetal cortex and adult non-cortical brain tissues (Figure 4). These findings are consistent with the hypothesis that development-stage specific eQTL patterns impact on disease-relevant mechanisms and thus may contribute to the proposed temporal phases of PD pathogenesis^47^.

**Figure 4:**
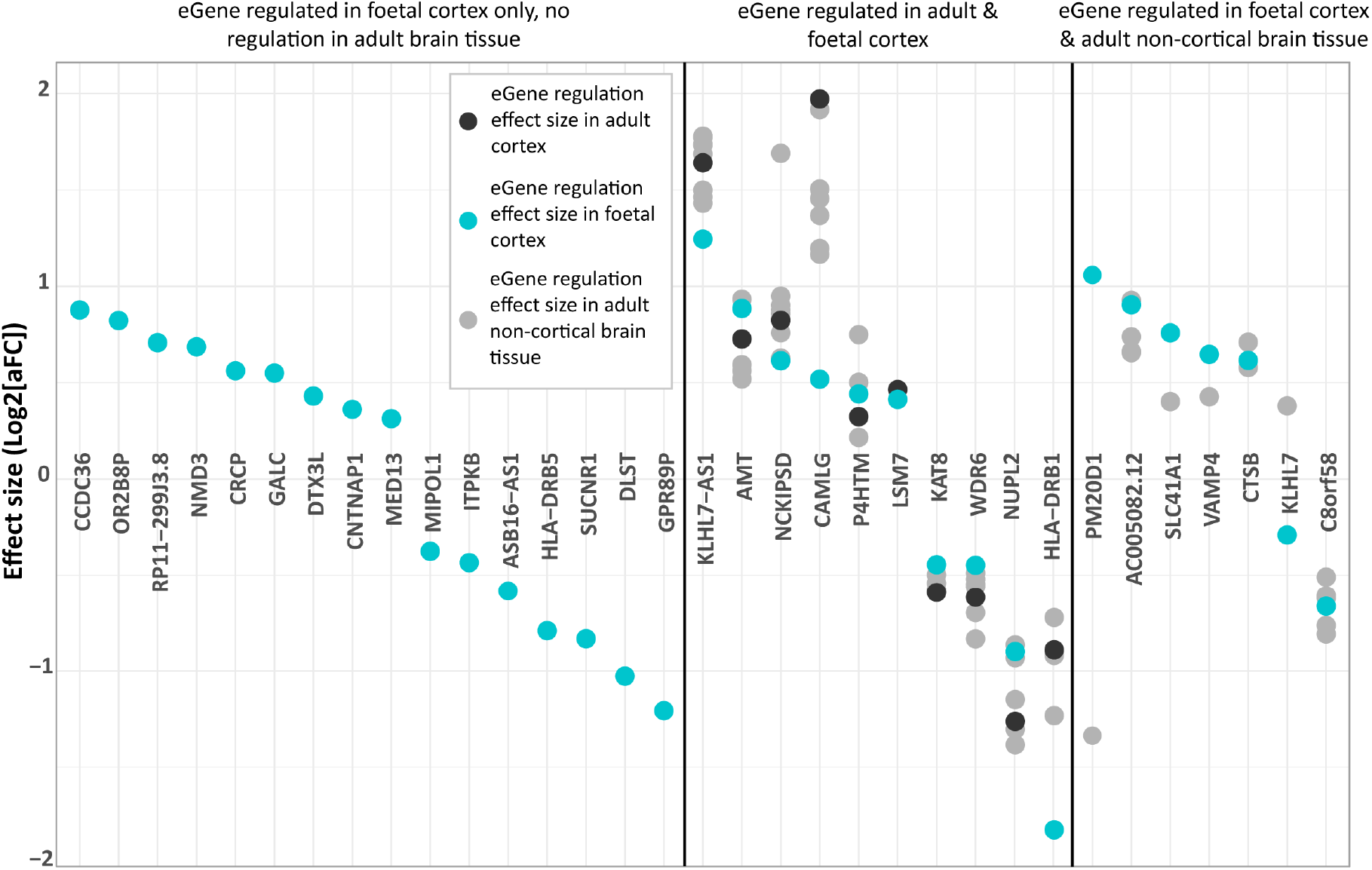
Gene regulation in the foetal cortex compared to the adult cortex. The leftmost section shows genes that are regulated only in the foetal cortex, with no eQTLs seen in any of the 13 adult brain tissues. The middle section shows genes that are regulated in both the foetal and adult cortex. The black dots show the regulation effect size of the gene in the adult cortex, and the grey dots show the regulation effect size of the gene across the different brain tissues (where an eQTL is seen). The rightmost section shows genes that are regulated in both the foetal cortex and adult non-cortical brain tissue.

### Louvain clustering highlights nine intra-connected protein clusters, enriched for disease-relevant, biological pathways

Network representations of complex datasets can aid the identification of biological relationships that are often not identified by enrichment analyses^48^. We used a Louvain clustering algorithm to identify clusters of interacting genes and proteins from within a Protein-Protein Interaction Network (PPIN) generated from the 523 eQTL regulated genes (518 adult tissue eQTLs and the 5 unique foetal cortex eQTLs; Supplementary table 9). Nine significant (p<0.05) clusters consisting of 122 genes were identified within the high-confidence PPIN (Figure 5). The genes within each cluster were regulated by between 5 and 18 PD-SNPs (Supplementary table 9) and every cluster contained at least two genes that were co-regulated by a single SNP (Supplementary figure 4). Notably, genes that were subject to *trans*-acting eQTLs were central to the definition and identification of several clusters (Figure 5). Pathway analysis (g:Profiler) of the genes within the individual clusters revealed enrichment in categories that included immunological surveillance (cluster 7), synaptic vesicle recycling (cluster 5) and microtubule polymerisation (cluster 3) (Supplementary table 10).

**Figure 5:**
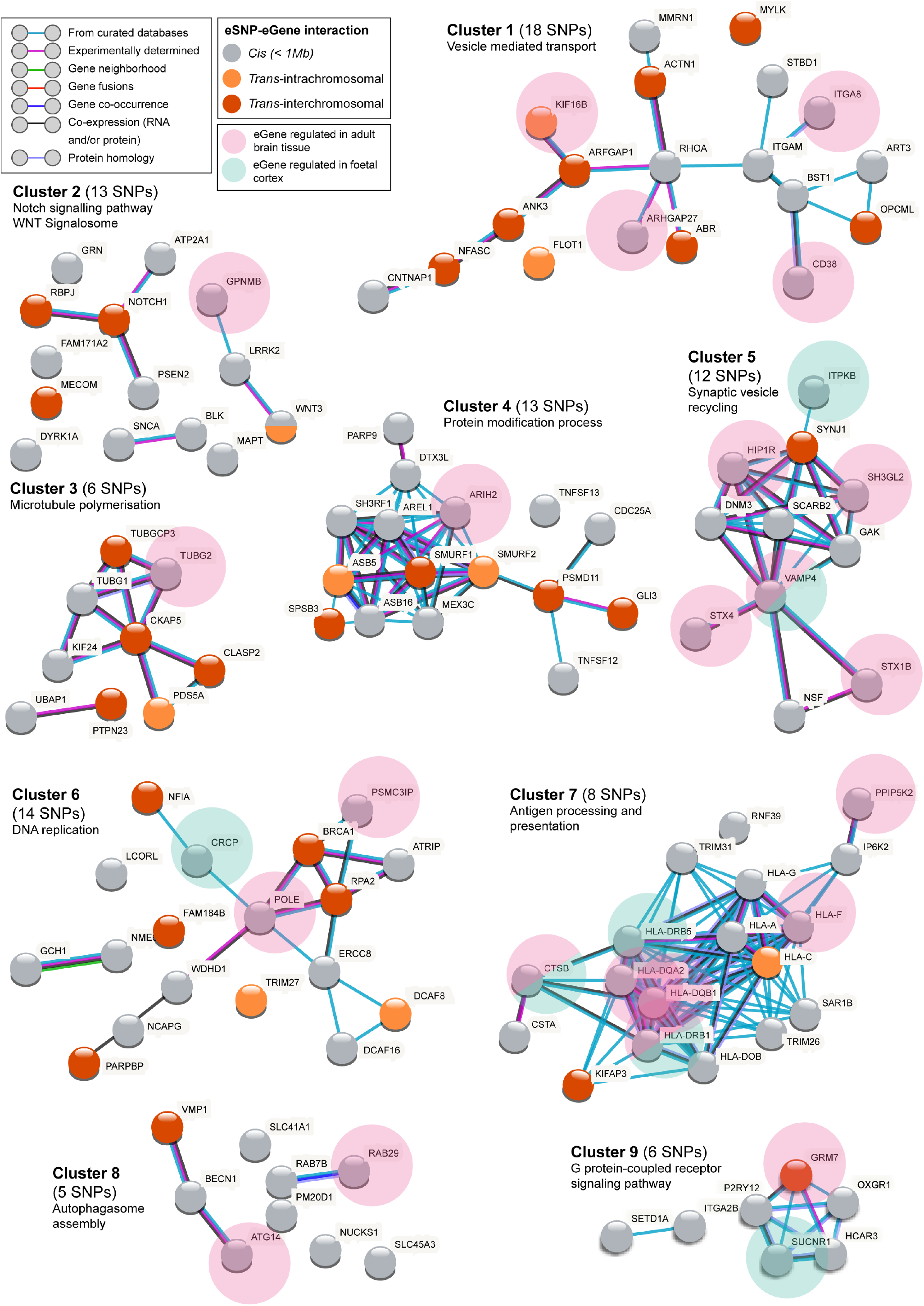
Louvain Clustering analysis highlights nine significant clusters, indicative of biological connectivity. The grey and orange shading of the nodes is indicative of whether the gene is subject to regulation via *cis*- or *trans*-mechanisms. The pink and turquoise shaded circles indicate genes that are regulated in adult brain tissue and foetal cortex respectively. STRING PPI confidence level = high (0.700); text-mining connections removed. The clusters were also analysed in STRING with an increased stringency (including only experimentally determined and curated database connections, confidence level: 0.700); however, this led to very few changes, with cluster 6 the only cluster to lose any connectivity within the cluster (WDHD1, NCAPG and PARPBP no longer connect). Experimentally determined: imported from experimental repositories; Gene neighbourhood: similar genomic context in different species suggest a similar function of the proteins; Gene fusions: fused proteins are recognised by orthology of the fused parts to other, non-fused proteins; Gene co-occurrence: indicates the presence of a specific gene pair is in agreement in all species – must be expressed together; Co-expression: predicted association between genes/proteins based on RNA and/or protein expression.

*Makarious* et al recently used a multi-modal machine learning approach, incorporating multi-omics datasets, to inform and improve predictions of PD^39^. Beyond the 90 GWAS SNP signals (which collectively were the top genetic feature), they also identified rs10835060 and rs4238361 as two SNPs that impact on PD biology. CoDeS3D analysis identified eQTLs for both rs10835060 (*KRTAP5-AS1, CCDC88A, KRTAP5-5, BRSK2*) and rs4238361 (*USP47*; and *RP11-507B12.2* and *RP11-259A24.1*) (Supplementary table 11). Notably, BRSK2 co-locates with cluster 2 through an established interaction with the Tau encoding *MAPT* gene^49^. The model also highlighted 29 genes through transcriptomic analysis. Three of these genes (*MMP9, TRIM4* and *SYS1*) integrate into clusters 1, 4 and 5, respectively (Supplementary figure 5). The co-location of genes and eQTLs, identified as being important for PD diagnosis^39^, within the nine clusters supports the potential importance of the gene-gene interactions and enriched pathways in PD.

402 genes did not segregate in the nine clusters. Of note, 211 of the 402 genes (52.5%) had not previously been associated with PD GWAS loci (iPDGC PD browser^40^). Of the 211, 123 genes were regulated by *trans*-eQTL-gene connections. Notably, iPDGC identified five genes (*DNAI1, EYA4, LYVE1, MYO5B, PDZRN4*) as being regulated by *cis*-eQTLs yet. These five genes were exclusively identified through *trans*-eQTL regulatory connections in our analysis.

## Discussion

Assigning functionality to PD-SNPs is a critical step towards determining how they contribute to the risk of PD development. In this study, we identified 518 genes whose expression was regulated in *cis* or *trans* by PD-SNPs, and the tissues where this regulation occurs. We also demonstrated that 22 PD-SNPs impact the regulation of a subset of 16 genes solely in the foetal cortex, and a further 10 genes in both the foetal and adult cortex. Of all 523 identified genes, a subset of 122 *cis*- and *trans*- regulated genes formed nine clusters within a protein:protein interaction network that were enriched for specific biological pathways, some of which have not been previously associated with PD. Our findings support the hypothesis that both *cis*- and *trans*- dysregulation of gene expression contributes to the risk of PD and provide insight into possible disease-causing mechanisms.

*SYNJ1* encodes synaptojanin-1, a presynaptic phosphoinositide phosphatase that dephosphorylates PI(4,5)P_2_ to trigger the removal of the clathrin coat during synaptic vesicle recycling^50^. *SYNJ1* is a highly constrained gene (LOEUF score = 0.33) and rare missense mutations in *SYNJ1* have been linked to early-onset Parkinsonism^51^. Despite this, GWA studies have not identified any SNPs proximal to *SYNJ1* as being significantly associated with PD, nor have they attributed any significant PD SNP (near or far) to *SYNJ1*. Critically, we identified that the PD-associated SNP rs10847864 acts as a *trans*-acting eQTL for *SYNJ1* expression. rs10847864 is intronic to, and also acts as a *cis*-eQTL with, *HIP1R*, another gene that is involved in clathrin mediated endocytosis^52–54^. Our discovery of the *trans*-eQTL-*SYNJ1* connection merges observations from population level (i.e. GWAS) and familial studies, reinforcing the potential importance of *SYNJ1* in PD.

Our analysis identified *trans*-eQTLs for approximately two-thirds of the known PD-SNPs. Of note, *RAI14* (retinoic acid induced 14) is regulated by two *trans*-eQTLs, involving two independent PD-SNPs (rs2251086, chr.15 and rs55818311, chr.19). Although not yet directly linked to PD, *RAI14* (and its encoded protein ankycorbin) has been shown to play a role in the inflammatory response in glial cells^55^, and in the establishment of neuronal morphology^56^; both of which are pathways of known importance in PD pathogenesis^57^. Retinoic acid, a regulator of *RAI14* (one of multiple roles of retinoic acid), is being explored as a potential therapeutic target for PD^58^.

Our results provide support for the role of peripheral tissues in PD, notably the oesophagus and thyroid. Firstly, the oesophagus is enriched for *cis* and *trans* regulatory eQTLs. rs76904798 (PD odds ratio (OR) = 1.155) is an eQTL that upregulates *LRRK2* expression in 19 peripheral tissues, including in the oesophagus. Notably, this *cis*-eQTL with *LRRK2* is not identified in any CNS tissues. Secondly, we identified the thyroid tissue as being enriched for eQTLs, many of which were not represented in CNS tissues. The thyroid is a component of the dopaminergic system and hypothalamic–pituitary–thyroid axis network^61^. A potential link between thyroid hormone disorders, PD risk, and symptom severity has been suggested^62^. Specifically, one study identified patients with hypothyroidism to have a two-fold elevated risk of developing PD^63^. Collectively, these findings support the growing body of evidence for the importance of the oesophagus^64,65^ and thyroid^62^ in PD.

We hypothesised that genes regulated by PD-SNPs in foetal cortical tissue may contribute to potential neurodevelopmental aspects of PD^24,45,46^. Sixteen genes were regulated by PD-SNP eQTLs within the foetal cortex. Two of these genes, *CNTNAP1* and *GALC* are particularly notable. *CNTNAP1* encodes Caspr1, a Neurexin family membrane protein. Reductions in Caspr1 concentrations delay cortical neuron and astrocyte formation in the mouse developing cerebral cortex^66^. *GALC* encodes a lysosomal galactosylceramidase that ensures normal turnover of myelin^67^ and has been linked to neuronal vulnerability^68^. While connections between the remaining 14 genes and PD development are less clear, we speculate that SNP mediated regulation of these genes specific to the foetal cortex may contribute to early neuro-developmental disturbances that render an individual more susceptible to PD.

Our analyses identified nine clusters that are enriched for specific biological processes and pathways, some of which have not previously been associated with PD. Dysregulated expression of the components within these pathways is potentially the basis of the risk conferred by the PD-SNPs. The clusters aid in understanding how PD SNPs mechanistically contribute to disease risk, and some interesting points can be drawn from these. For example, genes within cluster 6 are enriched for functions in DNA replication and repair, a pathway previously associated with the development of other neurodegenerative diseases^69^. Notably, *BRCA1* and *RPA2* (both previously linked to DNA damage response and repair^70,71^) are regulated in *trans* by PD-SNPs (rs11950533; rs9568188; rs62053943) and are central to cluster 6. It is notable that cluster 6 contains several factors associated with PARP1 activity (e.g. the PARP1 binding protein (PARPBP) and BRCA1) that link this cluster to the repair of SSBs which are enriched at neuronal enhancers in post-mitotic neurons^72^. A further example is the regulation of autophagy initiation by PD-SNPs, highlighted in cluster 8. Three interacting proteins within cluster 8, encoded by *VMP1, BECN1*, and *ATG14* are each regulated by a different PD-SNP (rs12951632 chr. 17; rs11158026 chr. 14; rs10748818 chr. 10; Supplementary table 3), including a *trans*-eQTL connection regulating *VMP1*. rs12951632 (OR= 0.93) and rs11158026 (OR= 0.919) are protective PD-SNPs that respectively increase *BECN1* and *ATG14* expression, two interacting core components of the PI3-kinase complex, required for autophagosome formation^73^. Individuals with both of these variants would potentially have increased autophagic capacity relative to individuals with one variant.

The genes in cluster 7 are strongly enriched for antigen processing and presentation – which is increasingly being implicated in the progression of PD^74^. Both rs504594 and rs9261484 are associated with a reduced risk of developing PD (OR = 0.8457 and OR = 0.9385 respectively). We identified a spatial eQTL between rs504594 and *HLA-DRB1* in both the foetal cortex and adult brain (including cortex and SN), implicating this regulatory eQTL-gene connection in both the neurodevelopment and neurodegenerative stages of PD. Interestingly, rs504594 (previous ID: rs112485776) was recently validated in a SNP-level meta-analysis (OR = 0.87), with results displaying no residual HLA effect in adults after adjusting for the SNP^75^. Instead, three amino acid polymorphisms within the *HLA-DRB1* gene were identified as drivers of the association between the HLA region and PD risk^75^. We agree that the impacts of rs504594 are contingent upon the *HLA-DRB1* allele - both in terms of regulation and protein sequence. We contend that the effects of the rs504594 eQTL are developmental. Future studies must untangle these developmental effects and identify the neurodevelopmental stages that may prime certain individuals to be more vulnerable to later triggering mechanisms. We note that such interpretation should be taken with caution given the highly polymorphic nature of the HLA-region.

We acknowledge several limitations to our analysis. Firstly, the Hi-C cohort and GTEx libraries were generated from unrelated samples that were not age or gender matched. Secondly, the sampled donors in GTEx are predominantly of European descent, limiting the significance of our findings to this ethnicity. However, the GWAS cohort also used individuals of European ancestry, meaning for this analysis the datasets were congruent. Thirdly, our eQTL analysis assumes that mRNA concentration correlates directly with protein levels. While it is true that protein levels are to some extent determined by their mRNA concentration, there are many post-transcriptional processes that can lead to a deviation from the expected correlation^76^. The fourth limitation is that eQTL data represent composite datasets across developmental periods (e.g. foetal samples were aged from 14-21 weeks post-conception and the adult samples were from individuals aged 21-70 years). Despite such limitations, the identification of *trans*-eQTLs is a particular strength of our methodology, relying on captured contacts within the genome organisation to reduce the impact of multiple testing correction. As such, we contend that these limitations do not invalidate the significance of our findings of *trans*-acting eQTLs and the genes that they impact.

## Conclusions

Understanding the functional impact of PD-SNPs is critical to our understanding of how these variants contribute to the development and clinical presentation of PD. Our functional interpretation of PD-associated SNPs integrates individual loci into a gene regulatory network, which includes genes with and without prior PD associations. The regulatory network includes clusters, and within them genes, that are enriched for biological functions that have known, putative or previously unknown roles in PD. Development-specific changes to this network (within the foetal cortex) are suggestive of roles for neurodevelopmental changes being early contributors to PD disease risk. Similarly, enrichments for regulatory changes within peripheral tissues may indicate a greater role for these tissues in PD than is currently appreciated. Collectively, our findings not only contribute to an overall understanding of the multiple biological pathways associated with PD risk loci, but also highlight the potential utility of gene regulatory networks when considering etiological subtypes of PD.

## Acknowledgements

SF, WS, TF, AC, and JOS were funded by the Michael J Fox Foundation for Parkinson’s research and the Silverstein Foundation for Parkinson’s with GBA – grant ID 16229 to JOS. SF and TF were funded by the Dines Family Charitable Trust. AC received grant funding from the Australian government. SG was funded by an MBIE Catalyst Grant (The New Zealand-Australia LifeCourse Collaboration on Genes, Environment, Nutrition and Obesity (GENO); UOAX1611). EG was funded by University of Auckland Doctoral Scholarship.

## Author Contributions

SF performed analyses and wrote the manuscript. TF and EG contributed to the preparation of eQTL and Hi-C datasets used in the manuscript. TF and WS contributed to the development of CoDeS3D. SG contributed to the network analysis script and the associated dataset. TF, EG, and SG all commented on the manuscript. WS and AC advised on the study and co-wrote the manuscript. JOS led the study and co-wrote the manuscript.

## Competing Interests Statement

The authors declare no competing interests

## Supplementary Material

Supplementary tables are available at:

https://github.com/sfar956/PDGWAS_regulatorynetworks

## Supplementary Figures

**Supplementary figure 1:**
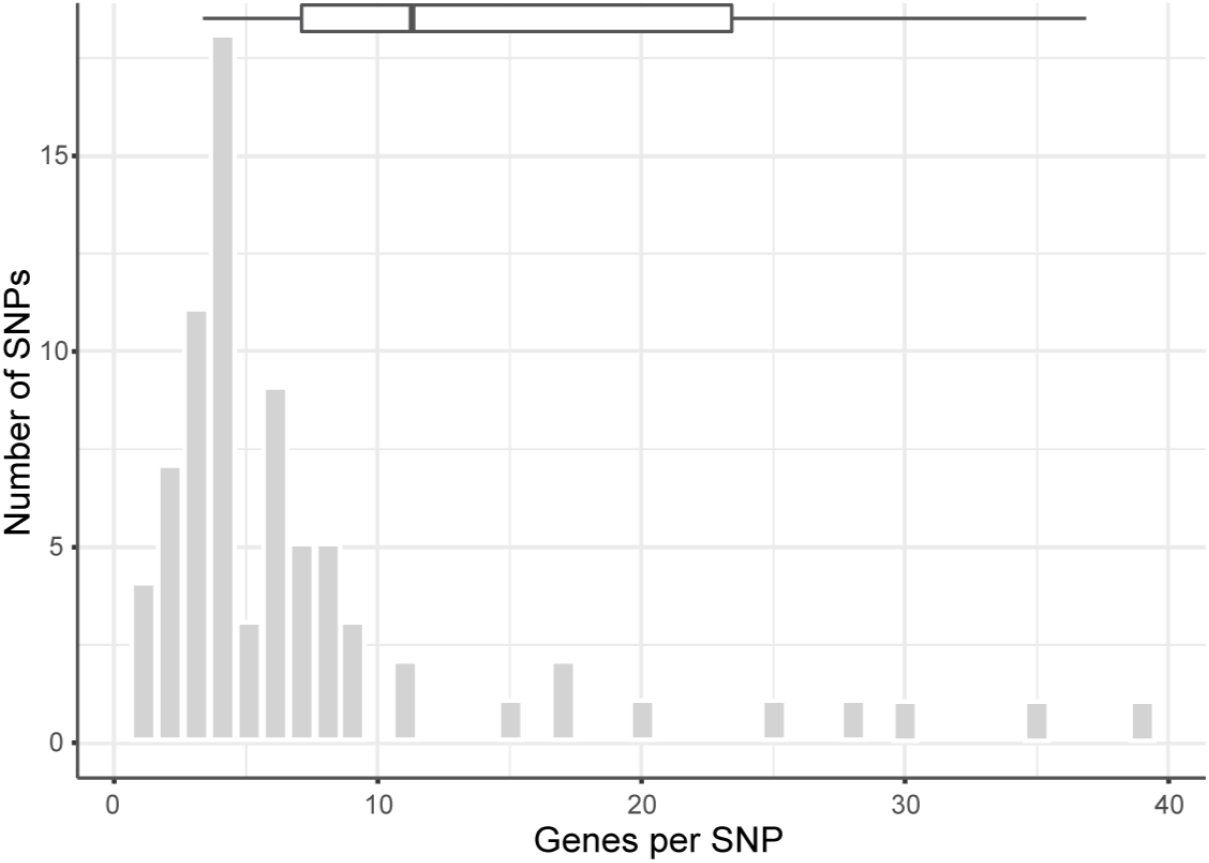
Number of genes regulated per SNP. Each of the 76 SNPs has an eQTL with between 1 and 39 genes. The boxplot represents the median and interquartile range.

**Supplementary figure 2:**
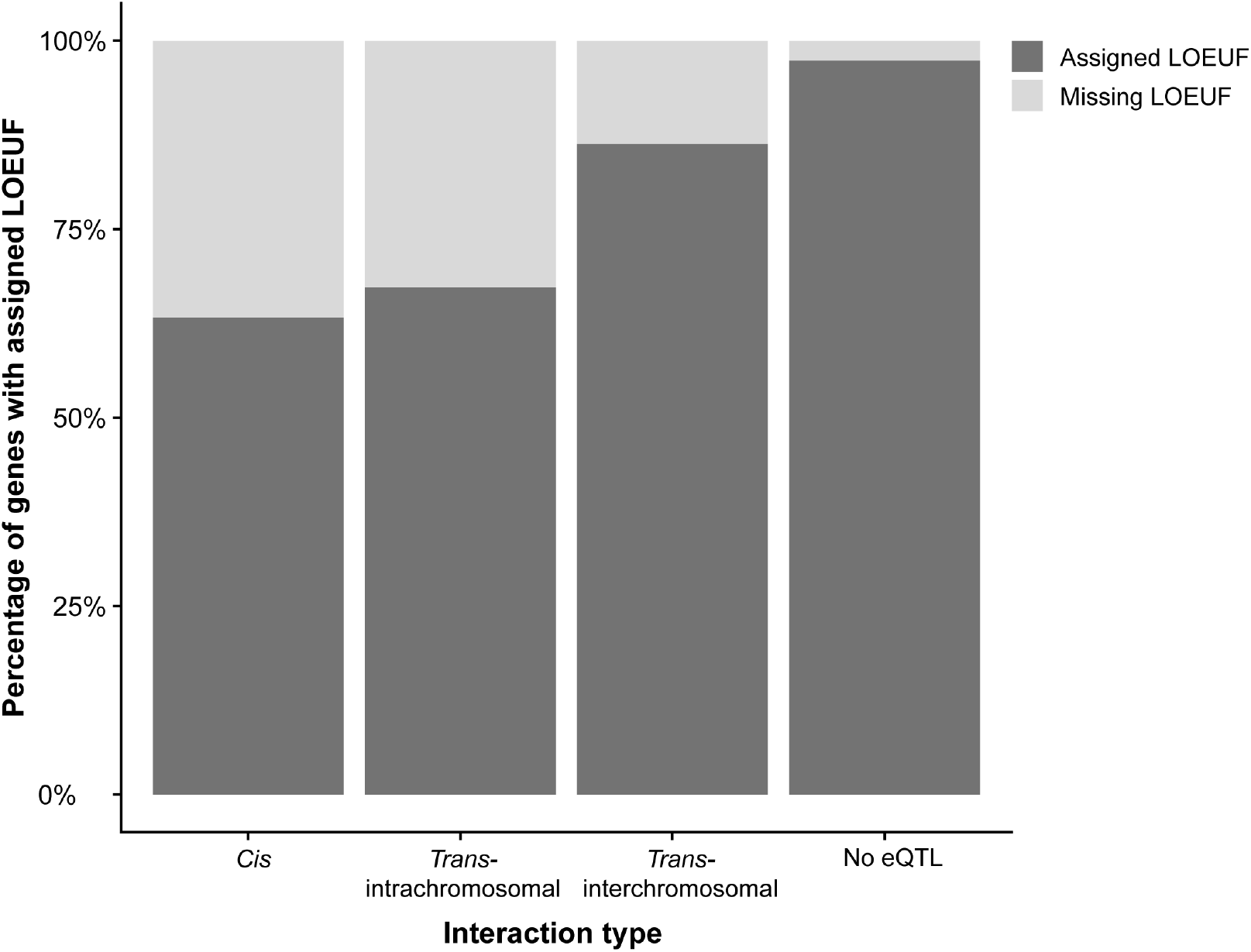
The proportion of genes that are loss of function intolerant increases as the eQTL distance increases. Not all genes have an assigned LOEUF score. The plot shows the percentage of genes that have no assigned score for each of the eQTL categories.

**Supplementary figure 3:**
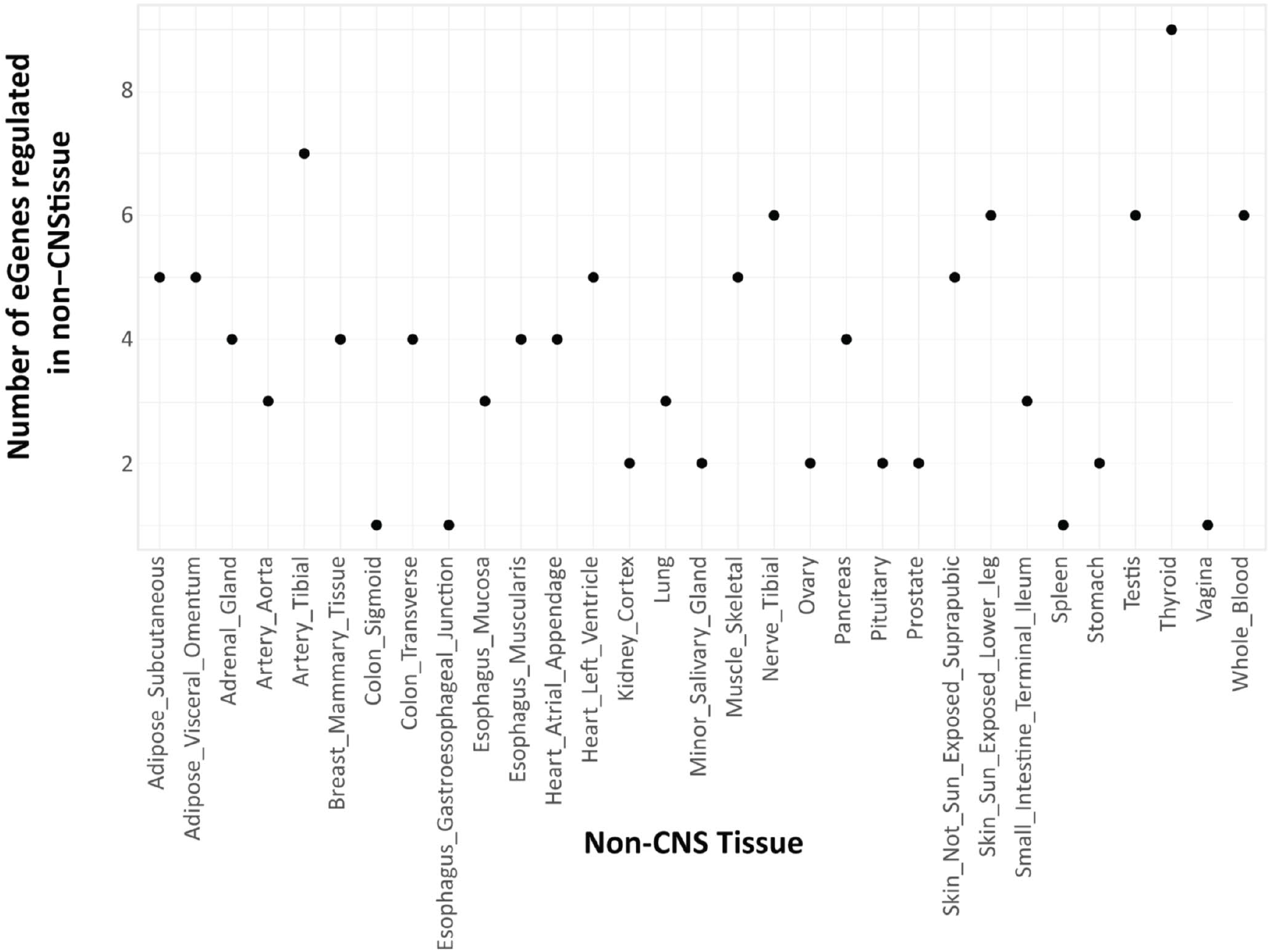
Genes subject to regulation in non-CNS tissues only. Our analysis identified a subset of 21 PD GWAS SNPs that regulate genes only within non-CNS tissues (*i.e*. these SNPs had no regulatory connections within brain tissues).

**Supplementary figure 4:**
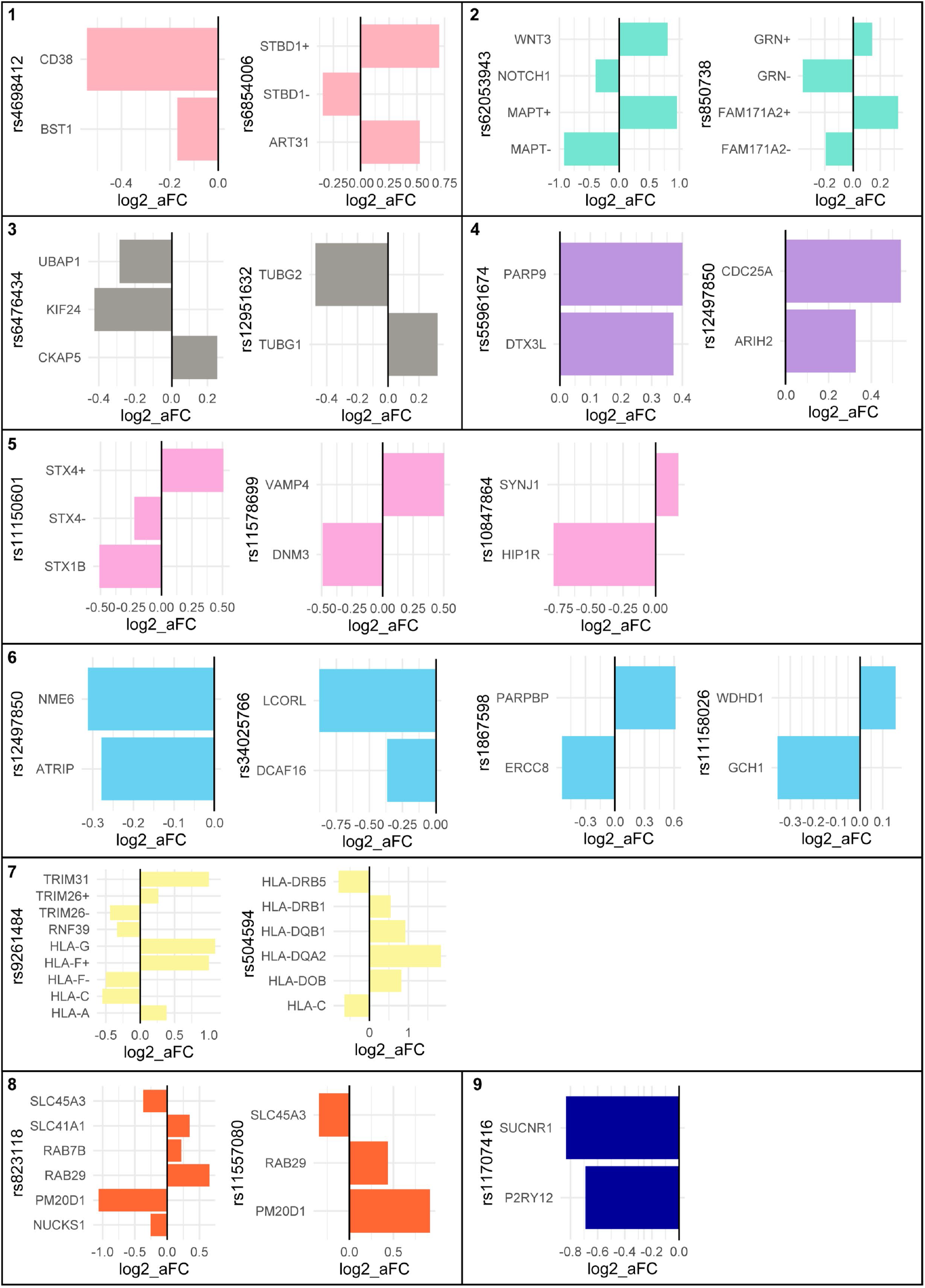
SNPs co-regulate multiple genes within individual clusters. For each of the seven clusters there is at least one SNP that co-regulates more than one of the genes within that cluster. The SNPs do not co-regulate the genes in the same direction in most instances. In some cases where the SNP regulates the expression of one gene in multiple tissues, the regulation may be positive in some tissues, but negative in others. For example expression of STBD1 is down-regulated in 7 tissues but up-regulated in the testis and lung.

**Supplementary figure 5:**
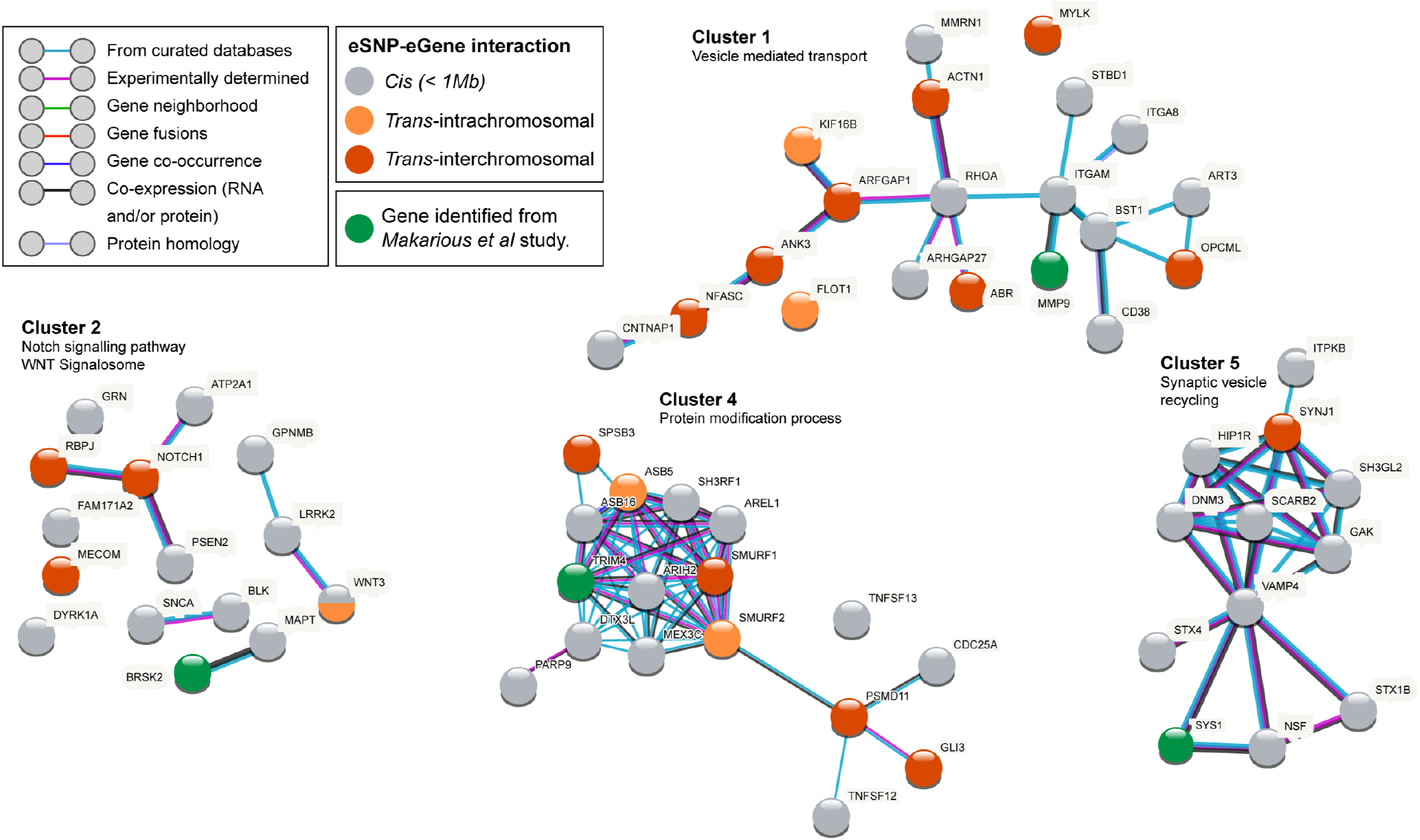
Genes and variants identified by *Makarious* et al connect into clusters 1, 2, 4 and 5. *Makarious* et al identified a set of genes and variants that affect the polygenic risk score for diagnosis of PD. A subset of these genes connect into the clusters identified through our analysis, showing a convergence between the two datasets, and further confirming the importance of the enriched pathways to PD biology. Adapted from main figure 5. The green shading of the nodes indicates genes that are identified from the *Makarious* et al dataset.

## Notes

https://github.com/sfar956/PDGWAS_regulatorynetworks

### Competing Interest Statement

The authors have declared no competing interest.

